# Disruption of the polyuracil tract can greatly enhance immunostimulatory properties of influenza delVGs

**DOI:** 10.64898/2026.10.13.758203

**Authors:** Sydney N.Z. Jordan, Marisa Mendes, Alistair B. Russell

## Abstract

Non-standard viral genomes (nsVGs) are replication-incompetent genomes within viral populations whose presence has been associated with an enhanced immune response. This association has garnered interest in their use as potential antivirals, vaccine adjuvants, or general immune-modulators. In influenza A virus (IAV), the predominant nsVG is a large deletion in one of the three polymerase segments, which results in deletion-containing viral genomes (delVGs). Work from our group has found that individual delVGs can be recovered that are capable of stimulating high levels of interferon, however we and others have not previously found any consistent means of predicting or modifying determinants of this stimulation. Using high-throughput genetic approaches, we find that specific mutations disrupting the polyuracil tract in IAV delVGs can enhance interferon production by up to two orders of magnitude. Such variants can also associate with improved rescue in artificial systems engineered to exclude infectious virus, and can greatly suppress the growth of replication-competent virus. These findings indicate the polyuracil tract is a key, and unexpected, target for improvement of immunostimulatory delVGs derived from IAV.

## Introduction

Initially thought to be an artifact of high MOI laboratory-grown viruses, non-standard viral genomes (nsVGs), which are a class of replication-incompetent particles, have received renewed interest due to their recognition as both a component of clinical infections, and their potential use as platforms for the development of antivirals and vaccine adjuvants.^1–6^

nsVGs, originally referred to as “defective-interfering particles” or DIPs, have been identified across several virus families.^7^ While the genetic basis of nsVGs differs between viruses, the defining characteristic of these particles is their ability to rise in frequency in the viral population under high MOI propagation despite themselves being unable to replicate. These properties, as well as their association with an increase in activation of host immune pathways, are considered potentially advantageous for use in novel therapeutic development.

Influenza A virus (IAV), the major causative agent of human flu, is a negative-sense, single-stranded RNA virus, consisting of eight viral RNA segments. nsVGs in IAV commonly appear within the three largest, polymerase-encoding RNA molecules (PB2, PB1, PA).^8–12^ Due to these large internal deletions, these viruses are often referred to as deletion-containing viral genomes (delVGs). IAV delVGs, as they do not encode a functional polymerase, are unable to replicate their negative-sense viral ribonucleoprotein (vRNP) to positive-sense complementary ribonucleoprotein (cRNP) intermediates in the absence of complementing virus.^13–15^ However, despite producing orders of magnitude less RNA than a fully infectious counterpart, they remain capable of independently inducing an interferon response in the absence of infectious virus.^16,17^

While there has been observed a general association of nsVG-rich stocks of virus with enhanced interferon production, validation of individual immunostimulatory nsVGs has been limited to a handful of viruses.^16–20^ In respiratory syncytial virus, for instance, it has been established that RNA structures present in “copy-back” genomes can serve as potent RIG-I agonists. For IAV, however, it is unclear why some delVGs have the capacity to exhibit high levels of immune stimulation and others do not, as well as how one could modulate the induction of a given delVG. Originally, it was thought that the reduced size of a delVG RNA molecule enabled it to serve as a better ligand for RIG-I.^21^ However, our group demonstrated that interferon induction does not have a direct relationship with delVG length, leaving the origin of immune stimulation by delVGs uncertain.^17^ Therefore even though our group has identified at least one highly stimulatory delVG, PB1_177:385_—a deletion in PB1 removing all but the first 177 and last 385 nucleotides of the vRNA—we do not understand how to modulate, and ideally enhance, its stimulatory effect.^16,17^

Here, using a comprehensive mutational analysis of PB1_177:385_ as well as other delVGs in PB2 and PA, we identify disruption of the polyuracil tract as a means of greatly increasing interferon induction. Characterizing disruptions to the polyuracil tract in delVGs derived from PB1 specifically, we find they not only enhance the host interferon response, but can improve both viral yield and the disruption of the growth of a wild-type competitor. We further find that such variants would be challenging to recover from natural sequence, and suggest that screens similar to ours could be beneficial in the development of nsVGs with desirable features for potential therapeutic use.

## Results

### An immunostimulatory delVG induces an interferon response in a minority of cells

To assess the properties of PB1_177:385_ as an immunostimulatory delVG, we performed single-cell RNAseq (scRNAseq) of A549 cells infected at an MOI 0.1 with either PB1_177:385_ or a matched, contemporaneously-generated, wild-type control at 10 hours post-infection (hpi, genome sequence in Supplementary Data 1). This time point lies largely within the first round of influenza replication, simplifying comparisons between replication-competent and -incompetent infections.^22^ Both datasets exhibit similar infection rates, with 826 (50.4%) and 954 (36.3%) cells in wild-type and PB1_177:385_ infections, respectively, meeting sample-specific thresholds for infection derived as described in methods (Fig. 1a). Matching our prior work, we identify more cells producing type I or III interferon transcripts (IFN^+^) above our detection threshold in the PB1_177:385_ infection than a wild-type control.

**Figure 1.**
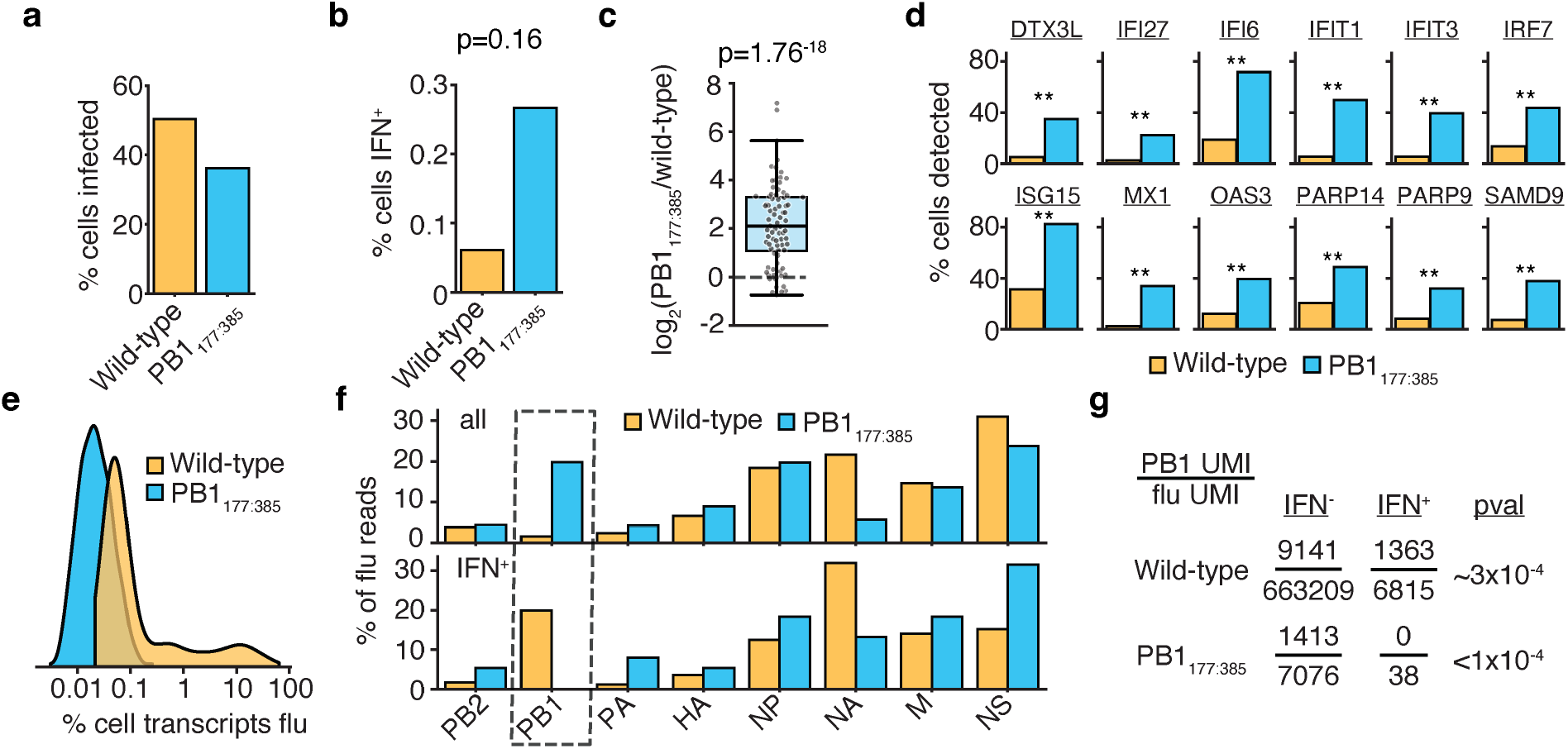
PB1_177:385_ induces interferon in a small number of cells, but evokes a strong ISG response. **a**,**b** Fraction of cells infected (a) or producing type I or III interferon (b) as defined using thresholds derived from spike-in of uninfected canine cells as presented in Supplementary Fig. 1. **c** Relative expression in pseudobulk of 88 ISGs defined from a previous study treating A549 cells with recombinant interferon as defined in Supplementary Data 3. Boxplots show mean, quartiles, and 1.5x inter-quartile range. **d** Percent of cells with detectable expression of the indicated gene in each dataset. Asterisks indicate significant difference, FDR <0.001. **e** Distribution of the fraction of reads in infected cells derived from influenza A virus. **f** Fraction of flu reads derived from each individual segment in each sample in pseudobulk of all infected cells or subsetted on just those producing interferon. PB1 highlighted for analysis in (g). **g** Analysis of frequency of PB1-derived reads in interferon-producing cells with an assumption that reads are drawn as a neutral process from a distribution represented by pseudobulk of all infected cells in each data set. p-values in (b, d, g) from Fisher’s exact test, values in (d) corrected using Benjamini-Hochberg. p-values in (c) one-sample t-test against the null hypothesis of a log ratio of 0.

However, while we observe an increase it only reflects a relatively modest number of cells producing detectable interferon transcripts; a single cell in the wild-type sample, versus seven in PB1_177:385_ (Fig. 1b). Although interestingly enough, the single cell in which we detect interferon in our wild-type infection appears to have been infected by a virion containing a PB1 delVG (Supplementary Data 2). As such, while the production of interferons appears to increase, it remains relatively rare, at least at this detection threshold at this time point. This increase fails to reach statistical significance, although this is likely due to the very low sampling in this dataset (p=0.16).

As interferons can induce a global antiviral state through paracrine signaling, we assessed whether the modest, not significant, increase in interferon-producing cells we observe in PB1_177:385_ instigated a much larger, and statistically robust, response throughout the host cell population. The antiviral state induced by interferons is mediated by the production of interferon-stimulated genes (ISGs). Using a set of ISGs defined from an RNAseq analysis of A549 cells treated with recombinant interferon, we assessed average expression of each ISG in pseudobulk analysis.^23^ Across a set of 88 ISGs we observe a median 4-fold increase in expression in PB1_177:385_ infection relative to wild-type, indicating that the PB1_177:385_ infection produced a global state consistent with enhanced interferon signaling (Fig. 1c). As the ISG response is generally thought to be bimodal (off/on), for 12 well-expressed ISGs where we observed an increase in PB1_177:385_ infection, we assessed whether these increases were the result of more cells expressing the indicated ISG relative to a wild-type infection.^24,25^ As anticipated, across all 12 ISGs there was a significant increase in cells expressing detectable transcripts from each individual ISG (Fig. 1d).

Having established that PB1_177:385_ induces a transcriptomic signature consistent with an interferon-driven antiviral state, we next sought to understand how the flu replication program may be altered by PB1_177:385_. As PB1_177:385_ is unable to generate new polymerase and engage in genome replication, flu transcripts in PB1_177:385_-infected cells are much lower on average than our matched, replication-competent, wild-type control (Fig 1e). This limits our analysis due to very sparse sampling of flu transcripts, and so rather than examine individual infected cells we decided to compare the flu transcriptional program in pseudobulk. Comparing both infections, we observe a two major alterations to the flu transcriptional program: 1) a relatively large increase in PB1 transcripts in PB1_177:385_ when compared against the other eight flu segments, and 2) a decrease in NA transcripts (Fig. 1f, top). We previously demonstrated that some delVGs can replicate more rapidly than full-length counterparts, however this is the first evidence that deletions may also enhance rates of transcription or transcript stability.^17^

Lastly, we explored whether variation in viral replication between PB1_177:385_-infected cells may explain the rare, interferon-positive events. We have previously leveraged viral heterogeneity to explore variation in the cellular response to flu, correlating reduced or absent transcription from the NS gene segment, which encodes the viral interferon antagonist NS1, and the presence of delVGs, with induction of an interferon response.^26^ Again, using pseudobulk analysis given the sparsity of reads from PB1_177:385_, we observe two features of note: 1) NS segment expression appears relatively unperturbed in interferon-producing cells, indicating NS1 antagonism was insufficient to prevent innate immune activation, and 2) there is a complete absence of transcripts from PB1_177:385_ itself. The latter was statistically significant, and odd both given the relative increase in PB1 transcripts associated with PB1_177:385_ as well as the fact that PB1_177:385_ itself seems to be responsible for the observed increases in interferon (Fig. 1f,g). From our further work, described below, we find that this absence may be explained by unsampled mutations in PB1_177:385_ that can greatly increase the interferon response.

### Disruption to specific sites in the influenza polyuracil tract can massively enhance the interferon response

As interferon induction remained relatively rare in cells infected by PB1_177:385_, it suggested that it may be possible to enhance or improve its capacity as an immunostimulatory delVG. Our prior work using length-variant libraries with combinatorial deletions and duplications failed to derive any fundamental principles of how to modify the immunostimulatory behavior of a given delVG, and so to assess the potential to improve or alter the behavior of PB1_177:385_ we turned to point mutation libraries generated by error-prone PCR.^17^

We already possessed such a library, as we successfully used this approach previously to identify and characterize sequences beyond the canonical U13/U12 promoter that are critical for replication in the presence of the flu export protein/polymerase cofactor NS2.^27–29^ To incorporate this library into a viral population, we performed influenza reverse genetics with the addition of a complementing, wild-type, virus. We then used these mixed populations to infect PB1-expressing A549 cells with an integrated *IFNL1* reporter, sorted cells by reporter expression, and sequenced variation in PB1_177:385_ in *IFNL1* -enriched and *IFNL1* -depleted populations.^16^ As we previously validated that this reporter correlates well with endogenous expression of both type I and III interferons we simplify our language in this section by referring to combined type I and type III responses as “IFN”, as in our single-cell work above.

Comparing variant abundance in IFN-enriched versus IFN-depleted populations, using both statistical and effect-size cutoffs, we identify several sites where variation co-associates with an enhanced interferon response (Fig. 2a,b). To our surprise, we do not identify any variants at the delVG junction that associate either positively (increased) or negatively (decreased) with an interferon response. This suggests that any aberrant RNA structure generated by the deletion itself has little impact on the observed interferon response to delVG IAV. For those sites where variation significantly associates with an enhanced interfon response, all are in the same functional region of the PB1 segment—the influenza polyuracil tract. This series of 5-6 uracils allows the flu polymerase to “stutter” and generate a polyA tail during transcription.^30,31^ Analysis of these positions suggests that, at several sites, introduction of a non-uracil base would be sufficient to enhance the interferon response to PB1_177:385_ (Fig. 2c). This effect appears to be specific to individual positions within the polyuracil tract, as not all sites in this tract exhibit strong selection in our analysis. Testing these predictions, we find that introducing a transversion mutation to the 20^th^ position (U20A) of PB1_177:385_ enhances the interferon response, whereas manipulating the 16^th^ position (U16A) does not in the PB1_177:385_ background (Fig. 2d).

**Figure 2.**
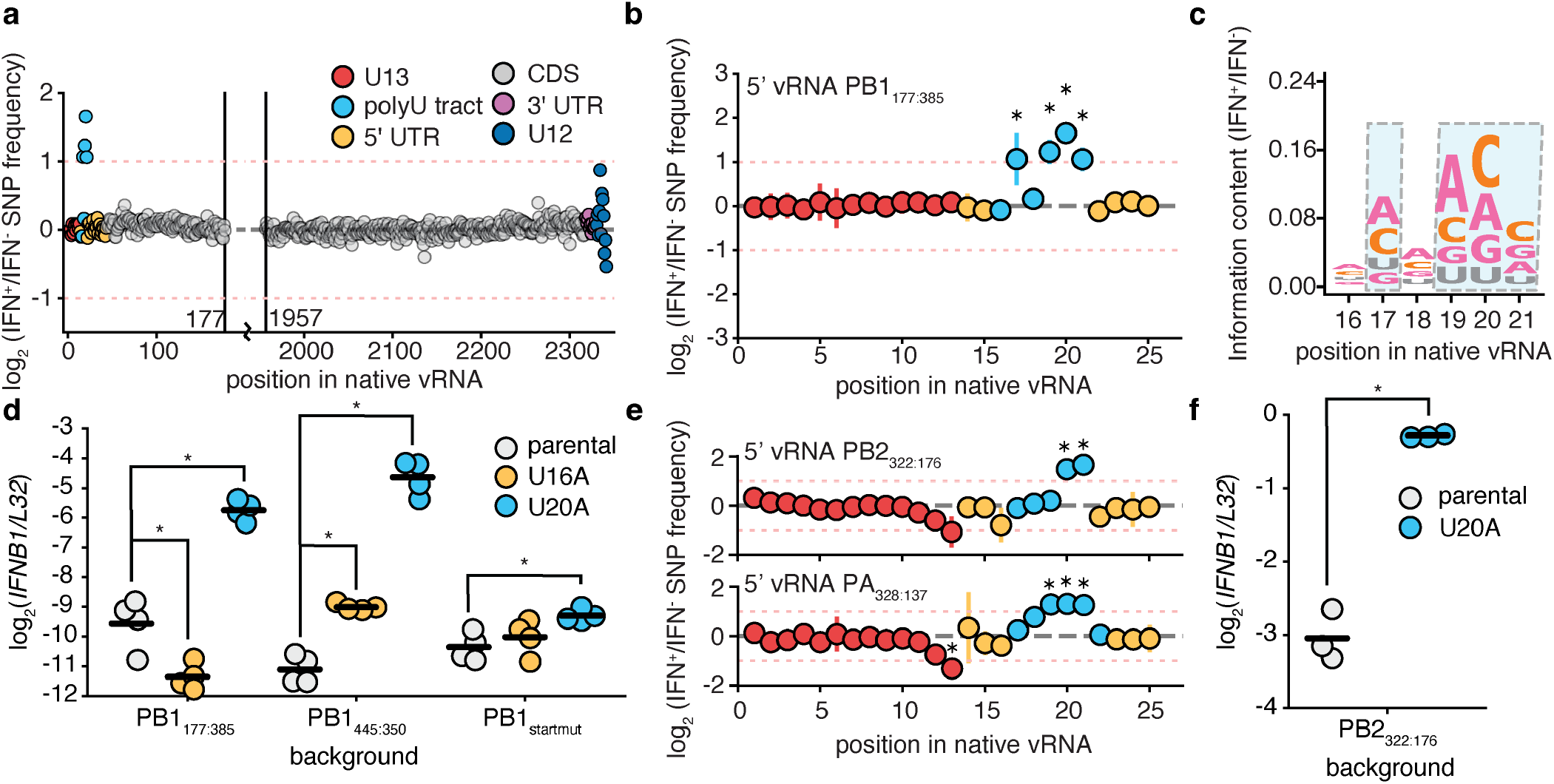
Mutational scanning identifies the polyuracil tract as a key determinant of interferon induction. **a** Mean ratio of non-wild-type nucleotide frequency at each position in PB1_177:385_ comparing IFN-enriched and -depleted A549 *IFNL1* -reporter cells, n=3. Dotted line indicates no difference in frequency between conditions, points above the line indicate mutations at that position associate with interfon induction, below, the opposite. Red dotted lines indicate two-fold differences in enrichment or depletion. Coordinates provided respective to full-length vRNA, regions of known functional significance noted. **b** The first 25 nucleotides of PB1_177:385_ analysed as in (a). Error bars are standard deviation, asterisks indicate sites with a greater than two-fold difference and significant enrichment or depletion relative to all sites measured by a two-tailed t-test with multiple-hypothesis correction by Benjamini-Hochberg at an FDR of 0.01. **c** Sequence logo of mean information content across all three replicates of each individual nucleotide in IFN-enriched versus IFN-depleted cells. Highlighted sites significant in (b). Data from sites 19-21 suggest any non-uracil nucleotide introduced at this position co-associates with interferon induction. **d** A549 cells were infected with the indicated variants at an MOI of 0.5 and *IFNB1* transcript levels relative to the housekeeping control *L32* were measured by qPCR performed on RNA harvested at 8 hpi. Asterisks indicate significant difference between a mutant and its parental background, two-tailed t-test with Benjamini-Hochberg correction, FDR 0.05, n=4. **e** Experiments with PB2_322:176_ and PA_328:137_ performed as in (a), analysis as in (b). **f** Experiment and analysis performed as in (d) with indicated variants, n=3. Original library mutation frequencies, distribution of variation with interferon, full datasets for PB2_322:176_ and PA_328:137_, and validation of IFN enrichment provided in Supplementary Fig. 2.

To determine if this effect is unique to PB1_177:385_, we introduced both the U20A and U16A mutations into additional backgrounds. For this we used both a less-stimulatory delVG (PB1_445:350_) and full-length PB1.^17^ As the latter could support replication, complicating an analysis of these positions, into this background we additionally introduced a mutation to the canonical start codon of PB1. In all templates U20A increased interferon transcription to a much greater degree than U16A, which either only slightly increased, or decreased, interferon transcription depending on the background (Fig. 2d). However, the magnitude of the increase produced by U20A was template dependent, with only a marginal increase in in *IFNB1* transcripts when U20A was introduced into full-length PB1 versus an almost 100-fold increase when introduced into PB1_445:350_.

While disruption to the polyuracil tract can be immunostimulatory in PB1, particularly when introduced into a delVG, we did not know whether this represents some unique feature of this RNA segment. We therefore used previously-identified delVGs in two other segments, PB2_322:176_ and PA_328:137_, generated libraries by error-prone PCR, and analyzed associations of mutations with interferon as with PB1_177:385_.^17^ Rather than sequence the entire template, we limited our analysis to the 5’ of vRNA/cRNA using RACE, to explore whether the polyuracil tract or other terminal nucleotides may play a role in innate immune induction in these backgrounds.^32^ As with PB1_177:385_, we predict that mutations to specific sites in the polyuracil tract increase the interferon response in both PB2 and PA templates (Fig. 2e). We confirmed this finding for PB2_322:176, U20A_ (Fig. 2f).

### Polyuracil tract mutations associate rarely with interferon in a high delVG population

Before exploring requirements for delVG PB1_U20A_ induction of an interferon response, we first sought to understand whether mutations to the polyuracil tract may be broadly responsible for the observation that IAV populations with high levels of delVGs induce a more potent immune response.^2,21,33,34^ While critical for polyadenylation of viral mRNAs, in delVGs we would not expect stringent selection for producing translation-competent mRNAs as they themselves do not encode any required proteins for replication. Indeed, quite the opposite, as proteins encoded by delVG can be inhibitory.^35^ Therefore mutations in this region may be found delVG genomes even if they would otherwise be purged from wild-type virus.

To address this question we resequenced the 5’ of vRNA and cRNA from PB2, PB1, and PA from a previously published high delVG infection. This experiment comprised two independent infections of A549 reporter cells expressing polymerase genes with a high delVG population followed by sorting on interferon expression.^26^ Our previous analysis of these data lack resolution of the polyuracil tract as it is at least partially overlapped by the most common set of sequencing primers used for IAV.^36^ These data allow us to determine whether polyuracil mutations may explain the increase in interferon during infection with a delVG-containing influenza population. Using a standard variant cutoff of 0.5%, we observe no detectable SNPs in either sample in the polyuracil tract of PB2, PB1, or PA (Supplementary Data 4).

We next lowered our variant calling threshold to 0.1% with a requirement that a variant is present in both replicates in either our IFN-enriched, or -depleted, populations to explore the potential contribution of rare variants to the interferon response. At this relaxed threshold we identify three potential SNPs in the polyuracil tracts of PB2 and PA (Fig 3a). Two of these, PA_U20C_ and PB2_U21C_, show evidence of co-enrichment with interferon, with PB2_U21C_ reaching significance even with only two replicates. The third, PA_U17C_, does not—matching predictions from our mutation libraries. This is consistent with a model where polymorphisms in the polyuracil tract can contribute in part to a delVG-specific interferon response, but only rarely and are unlikely to be the sole drivers of cellular detection of IAV. A caveat to this conclusion is that it is still formally possible that *de novo* variants arising during genome replication, too low to detect using our methods presented here, may be an undersampled and critical determinant of interferon induction in high delVG viral stocks.

**Figure 3.**
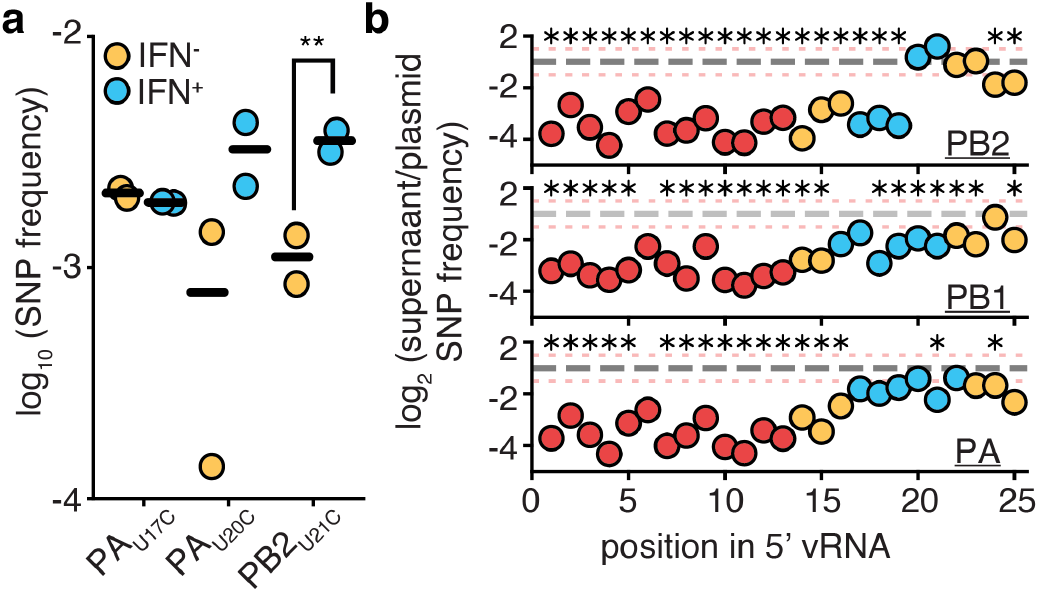
Polyuracil mutations are rare in a high delVG population and do not exhibit strong positive selection during delVG growth. **a** A high delVG population was previously used to infect A549 *IFNL1* reporter cells expressing each of the individual polymerase genes (PB2, PB1, or PA) at a genome-corrected MOI of 0.1 and sorted on reporter expression.^17^ PB2, PB1, and PA from two replicate infections were resequenced using 5’ RACE targeting vRNA and cRNA. Using a threshold of 0.1% variation/position with a requirement that both replicates identify the same SNP, we identify three potential SNPs in the polyuracil tract. Asterisks indicate significant difference btween IFN^+^ and IFN^-^, two-tailed t-test, p<0.05, n=2. Full data in Supplementary Data 4. **b** PB2, PB1, and PA libraries from Fig. 2 generated by reverse genetics were sequenced prior to interferon-enrichment experiments. Supernatant nucleotide frequencies were compared to original libraries (plasmid). Error bars lie within points presented, but would represent standard deviation. Asterisks represent points significantly different from no difference, one-sample t-test with Benjamini-Hochberg correction, FDR <0.1. Distribution of variation, full datasets for PB2_322:176_ and PA_328:137_ provided in Supplementary Fig. 3.

This leads us to ask, why are these variants so rare? As we sequenced supernatant vRNA from all three of our delVG mutation libraries after rescue by complementing virus, we could assess whether mutations to the polyuracil tract exhibit positive, negative, or neutral selection during viral growth and replication when introduced into a delVG. From these data, we find that polyuracil mutations generally exhibit either strongly negative or neutral selection, with small, not statistically-significant evidence of positive selection in a few sites in PB2 (Fig. 3b). The strong selection against mutations we see on PB1_U20_ is consistent with our prior work that mutations to this position decrease replication efficiency.^27^ Interestingly the few (rare) polyuracil polymorphisms we did observe are those we would have predicted to be under neutral or slightly positive selection. This is despite their association with interferon. From these observations we draw the conclusion that the the most stimulatory delVGs are, at best, rare in the population, as they lack any significant competitive advantage that could lead to their enrichment. Therefore the most stimulatory delVGs are not necessarily the best competitors nor the most aggressive, and efforts to identify them from natural diversity may be challenging owing to their rarity.

### Robust interferon induction by delVG PB1_U20A_ requires nuclear export, viral transcription, and RIG-I

It is not immediately clear why a mutation to the 20^th^ position of PB1 in a delVG could drive such a strong innate immune response. Moreover, this mutation can stimulate the transcription of interferons even in the absence of genome replication. Replication requries *de novo* polymerase synthesis—impossible for this virus in the absence of coinfection as it does not encode an intact polymerase complex.^13–15^ This means, either this mutation creates a vRNA that is such a potent innate immune agonist that a single molecule trafficing to the nucleus could be detected and drive a response, or, that disruption to the polyuracil tract has some secondary effect that is stimulating interferon production. To gain potential insight into the second possibility, we performed strand-specific total RNAseq on infections with PB1_445:350, U20A_ and PB1_445:350, U16A_. We opted to compare two different polyuracil mutations to to ameliorate any incidental differences that may occur through general manipulation of the polyuracil tract. We chose PB1_445:350_ as it is the template where we observed the largest increase in interferon transcription.

As expected, infection with PB1_445:350, U20A_ induces significant upregulation (40-60 fold) of type I and III interferons, as well as an upregulation of ISGs—indicating an active antiviral response, relative to PB1_445:350, U16A_ (Fig 4a). Focusing on differences in the virus itself, we first explored the relative amounts of each vRNA. We find no evidence of differences in total vRNA or the relative ratios of vRNA between U20A and U16A viruses. For both viruses PB1_445:350_ is somewhat under-represented in the vRNA pool, suggesting both PB1_445:350, U16A_ and PB1_445:350, U20A_ may have mild packaging defects (Fig. 4b). Next, exploring viral transcription we found differences in several segments. The most notable, when assessing both total numbers of transcripts and relative ratios of the viral mRNAs to one-another, was a significant decrease in mRNA from PB1_445:350, U20A_ relative to PB1_445:350, U16A_ (Fig. 4a,b). This is consistent with primer-extension experiments previously performed on PB1_U20A_ in minimal replication assays, which demonstrated mutations to this position decrease PB1 transcript abundance.^27^ No difference was observed in absolute transcripts derived from NS, with an increase in the fraction of total flu mRNAs from this segment in PB1_445:350, U20A_ due to relative decreases in mRNAs from the other segments. This again indicates that delVGs appear to be capable of inducing a robust interferon response even in the presence of NS1.

**Figure 4.**
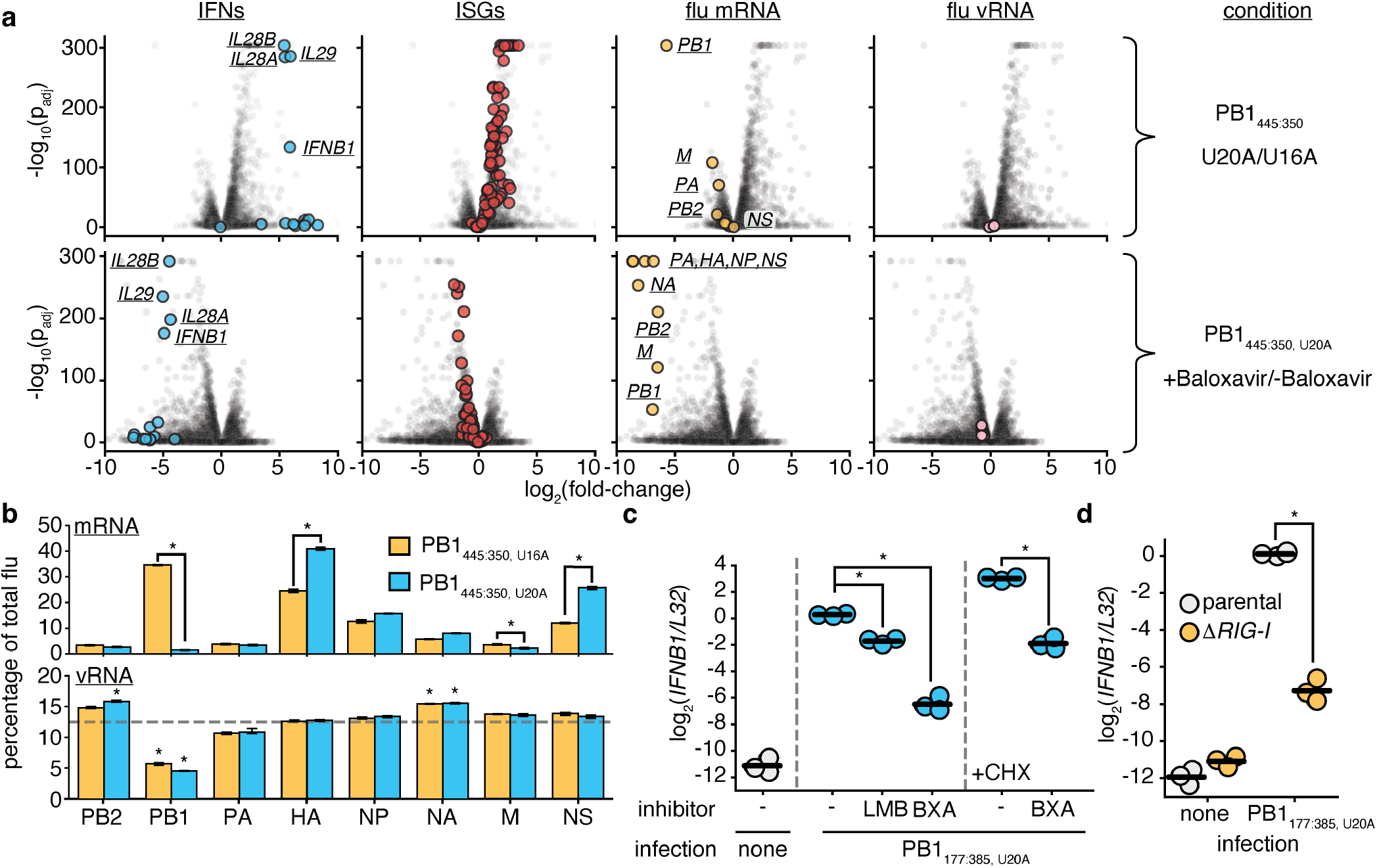
U20A mutation in PB1 delVG induces a potent interferon response via RIG-I requiring transcription and genome export. **a** Indicated variants were used to infect A549 cells at an MOI of 1 for 12h, RNA harvested and analysed by strand-specific total RNAseq after ribosomal RNA depletion. Strand-specific counts were generated by HTseq-count and indicated conditions compared by DeSeq2, n=3.^37,38^ IFN genes are all type I and type III genes, ISGs are defined as in Fig. 1c, flu mRNAs named if >2-fold effect size, or NS is highlighted due to its role in interferon suppression. **b** The relative fraction of length-corrected (+)-sense (top, mRNA) and (-)-sense (bottom, vRNA) reads from each segment were compared. For mRNA, asterisks represent differences between mutants, two-sample t-test with Benjamini-Hochberg correction FDR <0.001. For vRNA, dotted line represents equivalent frequency of each vRNA at 12.5% of flu reads (1/8), asterisks represent frequencies that differ from 12.5%, one-sample t-test with Benjamini-Hochberg correction FDR <0.01. **c** A549 cells were infected (or not) with PB1_177:385_ at an MOI of 2 and treated with baloxavir acid (BXA) or cycloheximide (CHX) at time of infection, or leptomycin B (LMB) 3 hpi to avoid interrupting early stages of the viral life cycle. RNA harvested 13hpi and analysed by qPCR to measure levels of *IFNB1* corrected for housekeeping control *L32*, n=3. Asterisks indicate significant reduction of *IFNB1* within panels against no leptomycin B or baloxavir, one-tailed t-test with Benjamini-Hochberg correction, FDR<0.05. Measurements of flu transcripts in Supplementary Fig. 4. **d** Indicated A549 cells were infected at an MOI of 0.5 with PB1_177:385_, RNA harvested at 14hpi analysed by qPCR to measure levels of *IFNB1* corrected for housekeeping control *L32*, n=3, data analysed as in (c).

Our group and others established that viral transcription can be a potent driver of the interferon response.^39–41^ However, here we observe a mutation that decreases mRNA abundance nevertheless associates with an incredibly strong interferon response. Does transcription remain critical to host detection of PB1_U20A_ in a delVG background? When we treat PB1_177:385, U20A_-infected cells with the viral transcriptional inhibitor baloxavir, we observe a significant decrease in the induction of *IFNB1* transcripts (Fig. 4c).^42^ Therefore despite the decrease in mRNA derived from this segment, transcription remains critical to detection of this virus. An additional, matched, stranded RNAseq dataset of PB1_445:350, U20A_confirmed that baloxavir suppresses positive strand synthesis in our infections as well as interferon and ISG induction, and indicates our findings are not unique to the PB1_177:385_ background (Fig. 4a).

In addition to a requirement for viral transcription, cellular detection of influenza requires active export from the nucleus via the CRM1 pathway, and does not absolutely require viral protein synthesis.^26,39,43^ To explore the relevance of these findings to delVG PB1_177:385, U20A_, we blocked CRM1-dependent nuclear export with leptomycin B, and explored whether viral transcription could still drive an interferon response in the presence of the translation inhibitor cycloheximide. Consistent with prior work, interferon transcription was reduced by leptomycin B and remained sensitive to baloxavir treatment even during translational block by cycloheximide (Fig. 4c). Lastly, the main pathway through which flu triggers an interferon response relies on the cytoplasmic sensor of triphosphorylated dsRNA, RIG-I.^44^ Using a knockout line previously generated and characterized by our group, we confirm that PB1_177:385, U20A_ induces a significantly reduced interferon response in the absence of RIG-I (Fig. 4d).^40^

Thus PB1_U20A_ in delVG backgrounds appears to generate some *de novo* RIG-I ligand, requiring flu transcription and active nuclear export, despite decreasing mRNA abundance from PB1. It seems unlikely that standard flu transcription products, appropriately capped and polyadenylated, are driving the response we see to PB1_U20A_, and that an alternate ligand, possibly the recently described ccRNAs, is the relevant target of RIG-I.^41,45^ Unfortunately in our experiments we capture (+)-sense RNA sequence adjacent to the polyuracil tract in the PB1_445:350, U20A_ infection in only a single read out of 100 million total paired-end reads, leaving the polyadenylation state, or lack thereof, of the reduced (+)-sense RNA species uncertain. While this single read was consistent with a ccRNA, we lack sufficient capture of this region to make strong statements. Conversely, we do capture polyadenylated mRNAs from PB1_445:350, U16A_, demonstrating a shortening of the polyuracil tract does not appear to completely inhibit polyadenylation (Supplementary Fig. 5).

### A U20A mutation in PB1 can enhance desirable features of candidate antiviral delVGs

Finally, we wished to assess two critical features of PB1_U20A_ to determine whether our work has identified a potential platform as a stimulatory delVG. The first is the capacity to grow this virus. Our prior work with PB1_177:385_ struggled due to a difficulty in rescuing this virus by reverse genetics, with titers around the limit of detection that needed subsequent passaging to achieve reasonable titers for experiments. At the time, we presumed this was due to innate immune induction leading to difficulties in rescue. However, to our surprise, we have consistently rescued improved titers of PB1_177:385, U20A_, in addition to its less stimulatory counterpart, PB1_177:385, U16A_ (Fig. 5a). As measuring infectious titers of an innate-immune stimulatory delVG may be inaccurate, we measured the total supernatant copies of the PA genome segment in reverse genetics rescues of U20A variants as a proxy for productive release of virions. Consistent with our titer observations, U20A greatly increased release of PA in either delVG background, although curiously not in full-length PB1 (Fig. 5b). We confirmed that introduction of this mutation into full-length PB1 did indeed result in decreases in titer as measured on PB1-complementing cells (Supplementary Fig. 6). Therefore U20A can improve rescue of delVG particles in reverse genetics, although in a template-specific fashion, despite itself reducing genome replication as demonstrated in our prior work.^27^ Given observations that delVGs can exhibit enhanced replication (although through complex mechanisms and not simply the shortest win, as observed by ourselves and others), we speculate that the U20A mutation may bring delVG replication “back in balance”, whereas when introduced into a full-length template it may disrupt pre-existing replication balance.^8,17,27,46–49^ Therefore while PB1_U20A_ may be at a disadvantage in competition with other variants within a single infected cell, when grown in single in a delVG it improves titer. These data may also explain why we see that PB1_445:350, U20A_ appears relatively stable at least through initial reverse genetics rescue, as we observe only a single read across all three replicates that does not support the U20A point mutation (Supplementary Fig. 7).

**Figure 5.**
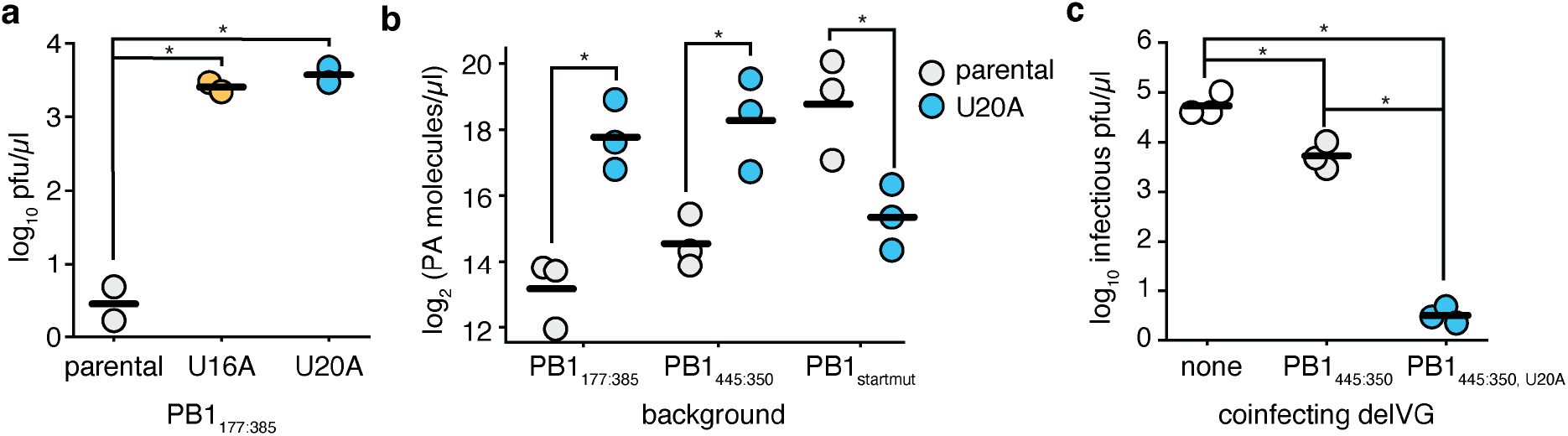
The PB1 U20A mutation improves delVG yield and antiviral activity. **a** Titers of indicated PB1_177:385_ variants after reverse genetics recovery as calculated on MDCK-SIAT1 cells expressing PB1. Asterisks indicate significant difference from parental, two-tailed t-test with Benjamini-Hochberg correction, FDR 0.05, n=2. **b** PA molecules from supernatant from experiments as in (a). Asterisks indicate significant difference from parental, two-tailed t-test with Benjamini-Hochberg correction, FDR 0.05, n=3. Titer data from rescue of full-length PB1 with a start codon mutant analogous to (a) presented in Supplementary Fig. 6. **c** A549 cells were infected at an MOI of 0.1 for 2h with the indicated variants before being challenged with wild-type influenza at an MOI of 0.01. Infections were allowed to proceed for 72h prior to supernatant harvest and assessment of growth of replication-competent virus. Asterisks indicate significantly different titers, ANOVA p<0.05 with post-hoc Tukey’s test FWER < 0.05.

The other desirable feature of an antiviral delVG platform is to inhibit the growth of a competitor virus. It is predicted that the capacity to stimulate an innate immune response would have a significantly stronger antiviral effect than simply directly competing within an infected cell.^33,49,50^ We compared the ability of PB1_445:350_ and PB1_445:350, U20A_ to act as antivirals in co-infection experiments with wild-type flu. We chose this template as the one in which we see the greatest differential in interferon, as in our total RNAseq experiment. After 72h of coinfection, allowing multi-cycle replication, we find that while PB1_445:350_ was able to decrease infectious virus by 1-log, PB1_445:350, U20A_ reduced growth of replication-competent virus by an incredible *∼* 4-logs (Fig. 5c). For influenza this phenotype is, to the best of our knowledge, only rivaled by the “OP7” virus, an interesting IAV strain with 37 point mutations in the M segment which also induces a potent interferon response.^5,51–53^

## Discussion

Through mutational scanning we have found that point mutations to the polyuracil tract can generate highly antiviral, immunostimulatory, influenza A virus delVGs, improving interferon induction by up to two logs and increasing antiviral effects by approximately three logs. It was somewhat surprising that a single mutation could have such a dramatic effect, rivaling mutational inactivation of major interferon antagonist NS1.^54^ Serendipitously, the mutation we focus on here, delVG PB1_U20A_, not only increased the interferon response, but also greatly improved our ability to recover this variant by reverse genetics, making it a prime candidate for further study.

Unexpectedly we did not identify any mutations around the delVG junction in PB1_177:385_ predicted to influence the interferon response. We still do not fully understand why this delVG co-associates with interferon, but given our difficulty in rescuing PB1_177:385_, ease of rescuing PB1_177:385, U20A_ and the absence of transcripts from PB1_177:385_ in interferon-positive cells in our scRNAseq dataset, it is tempting to speculate that we inadvertently enriched for polyuracil mutations during reverse genetics. Notably, we did still identify a large number of polyadenylated transcripts from PB1_177:385_ in our scRNAseq dataset, suggesting if U20A or similar mutations are present, they still remained rare. Moreover our analysis of a high delVG stock suggests that even if PB1_177:385_ is explained by enrichment of polyuracil mutations, these mutations are unlikely to globally explain why delVGs can be immunostimulatory. Further work will be necessary to provide a comprehensive explanation of their association with an interferon response.^2^ A key clue we may draw from our work here, however, is that a polyuracil mutation only drove a strong interferon response in a delVG background, indicating there must be some difference between these species and a full-length counterpart in terms of either generation of a RIG-I ligand or affinity of that ligand for RIG-I.

We were also initially surprised that interferon induction and competitive fitness were not particularly well-correlated.

This indicates it may be challenging to identify optimal antiviral candidates in naturally-occurring delVGs. Consistent with this logic, if immunostimulatory delVGs were highly fit then in natural IAV infections they would doubtless rise in frequency and collapse population fitness, potentially driving the virus to extinction. Therefore we would expect them to not be a common component of IAV infections, simply because they are incompatible with successful infection due to the sensitivity of influenza to interferon signaling.^54–58^ We speculate that, as with our prior work on the co-encoding of NS1 and NS2, and the work of ourselves and others on selection against non-polymerase delVGs, both the genomic architecture of IAV and selection pressures during genome replication may serve to limit the success of dangerous, interferon-stimulatory, variants in the population.^17,26,59^

Next, while the flu genome was originally thought to be the sole RIG-I ligand during infection, our findings here, our previous work, and the work of others demonstrate this is unlikely to be strictly true.^21,39–41,44,45,60,61^ This is unsurprising for a delVG in a polymerase segment, as if the genome itself were the sole ligand it would have to trigger a response when delivered as a single RNA molecule during its limited transit time in the cytoplasm towards the nucleus. While we did not fully characterize how mutations to the polyuracil tract drive such a strong interferon response, we can say with confidence that it relies on transcription and active nuclear export, and proceeds through the major pattern-recognition receptor for IAV, RIG-I. All of this is consistent with a body of work demonstrating that transcription products can be a source of RIG-I activation in IAV inefection.^39,40,45^ These data are also highly consistent work on a class of non-polyadenylated, capped, RNAs generated by flu called ccRNAs, which can, through association with small-viral RNAs (svRNAs) drive a strong interferon response.^41^ We anticipate modulation of ccRNA production as a likely explanation for our phenotypes.

Lastly, we believe that approaches such as ours will hopefully provide the raw material for use of delVGs, and other nsVGs, as therapeutics. The high level of intereferon induction we observe here is appealing for a vaccine adjuvant, as interferon signaling is considered critical for the production of a durable vaccine response.^62–65^ Whether higher is necessarily better will require further investigation, but with this work we now provide additional candidates with which to test these hypotheses.

Beyond the identification of immunostimulatory delVGs our work demonstrates the power of comprehensive mutational profiling and deliberate selection on desirable features. We would not have anticipated prior to our research that individual point mutations can cause such dramatic, and hopefully useful, impacts on delVGs, improving titers, interferon production, and antiviral responses. We hope others leverage the tools and approaches we provide here to develop nsVGs for eventual use in improving human health and combating viral disease.

## Methods

### Cell lines and viruses

The following cell lines were used in this study: HEK293T (ATCC CRL-3216), MDCK-SIAT1 (variant of the Madin-Darby canine kidney cell line overexpressing SIAT1 (Sigma-Aldrich 05071502)), MDCK-SIAT1 PB1, MDCK-SIAT1 PB2, A549 (human lung epithelial carcinoma cell line, ATCC CCL-185), and A549 pIFNL1-LNGFCR-p2A-zsGreen.^16,35,66^ Cell lines were tested for mycoplasma using the Invivogen Mycoplasma Detection Kit. All cell lines were maintained in D10 media (DMEM supplemented with 10% heat-inactivated fetal bovine serum and 2mM L-Glutamine) in a 37°C incubator with 5% CO_2_. Small molecules/drugs were used when indicated at the following concentrations, baloxavir acid, 100nM, cycloheximide, 50*µ*g/ml, leptomycin B, 10nM. The influenza strain A/WSN/1933 was used for all experiments. Genome sequence of A/WSN/1933, PB1_177:385_, PB1_445:350_, PB2_322:176_, and PA_328:137_ provided in Supplementary Data 1.

Wild-type A/WSN/1933 (H1N1) virus was created by reverse genetics using the bidirectional plasmids pHW181-PB2, pHW182-PB1, pHW183-PA, pHW184-HA, pHW185-NP, pHW186-NA, pHW187-M, and pHW188-NS.^67^ HEK293T and MDCK-SIAT1 cells were seeded in an 8:1 coculture and transfected using BioT (Bioland Scientific, LLC, #B0101) 24 hours later with equimolar reverse genetics plasmids. 24 hours post-transfection, D10 media was changed to Influenza Growth Medium (IGM, Opti-MEM supplemented with 0.04% bovine serum albumin fraction V, 100 *µ*g/ml of CaCl_2_, and 0.01% heat-inactivated fetal bovine serum). 48 hours post-transfection, viral supernatant was collected, centrifuged at 300g for 4 minutes to remove cellular debris, and aliquoted into cryovials to be stored at -80^*°*^C. Thawed aliquots were titered by TCID50 on MDCK-SIAT1 cells and calculated using the Reed and Muench formula.^68^ To create wild-type viral stocks with a low defective content, MDCK-SIAT1 cells were infected at an MOI of 0.01 and harvested 30 hours post-infection and titered.

Stocks of delVG virus for PB1_177:385_, PB1_445:350_, PB2_322:176_, PB1_startmut_ and associated polyuracil mutants were generated similary to wild-type A/WSN/1933 stocks with a few key differences: 1) complementing MDCK-SIAT1 cells were used to support growth of the delVG, 2) additional protein expression constructs for PB1 or PB2 were added to the initial transfection, 3) virus was harvested 72h post media-change, 4) titering was performed on complementing cells, and 5) for our start mutant plasmids we used the unidirectional rescue plasmid pHH21.^69^ For PB1_177:385_, PB2_322:176_, and PB2_322:176, U20A_ due to low levels of rescue from reverse genetics with viral titers close to, or at, the limit of detection, an additional 72h blind passage on complementing cells was necessary to procure sufficient material for experiments in Figs. 1 and 2.

### Reverse transcription and qPCR

For all experiments, RNA was purified using the Monarch Total RNA Miniprep Kit (New England Biolabs, #T2010), or the Quick-RNA Miniprep Kit (Zymo Research Corp, Cat #R1055). All experiments within the same panel, or sequencing experiments compared within this manuscript, used the same purification kit. For all qPCR experiments, cDNA was generated using the High Capacity First Strand Synthesis Kit (Applied Biosystems, 4368814). Reverse transcription was performed using the manufacturer’s protocol, except for our measurements of supernatant genome copy we used the universal vRNA primers as described in Hoffmann *et al*.^36^ Reaction conditions were an initial step of 10 minutes at 25^*°*^C, reverse transcription at 37^*°*^C for two hours, and an 85^*°*^C heat inactivation step for five minutes. All qPCR experiments used Luna Universal qPCR Master Mix (NEB #M3003), with primers specified in Supplementary Data 6. qPCR conditions for all experiments were as follows: denaturation at 95^*°*^C for one minute, then cycles of denaturation at 95^*°*^C for 15 seconds, and annealing and extension at 60^*°*^C fo 30 seconds. A melting curve was then performed to ensure all products are mono-peaked. For measurements of cell-free viral populations, 100 *µ*l of supernatant and 700 *µ*l of lysis buffer were combined and processed. For calculations of PA molecule numbers, values were compared against a molecular standard generated from PA-encoding plasmid DNA.

### Random mutagenesis of delVGs

Construction of the PB1_177:385_ library, and sequencing, was described previously.^27^ To construct PB2_322:176_ and PA_328:137_ libraries, we used primers flanking each template in a bidirectional reverse genetics plasmid and amplified the indicated delVG using the GeneMorph II Random Mutagenesis kit (Agilent, #200550) with a 54^*°*^C annealing temperature, 30 cycles, and a target template input matching manufacturer’s recommendations targeting a mutation rate of 1%. Products were isolated by gel electrophoresis, column-purified, subjected to 10 rounds of amplification using Q5 Hot Start High-Fidelity 2X Master Mix (New England Biolabs #M0494) to achieve sufficient concentration for library production, and cleaned with SPRI beads (Promega ProNex Size-Selective Purification System, #NG2001) at a 3:1 ratio. Vector backbones were prepared by inverse PCR using Q5 Hot Start High-Fidelity 2X Master Mix (New England Biolabs #M0494), gel purified, and cleaned with SPRI beads at a 3:1 ratio. Libraries were then assembled by Gibson using New England Biolabs NEBuilder HiFi (DNA Assembly Master Mix #E2621) at a 2:1 molar ratio of insert to vector. Assemblies were SPRI bead purified at a 1:1 ratio and electroporated into ElectroMax DH5*α* (ThermoScientific #11319019). All libraries were generated in triplicate, starting with three independent mutagenesis PCRs.

To rescue viral libraries for both supernatant and interferon measurements, we transfected an 8:1 coculture of MDCK-SIAT1 and 293T cells with helper plasmids encoding only the mRNA for A/WSN/1933 PB2, PB1, PA, NP, and NS2 and the library in a 1:1 ratio using BioT (Bioland Scientific LLC, # B1011) transfection reagent according to the manufacturer’s protocol. 24h post-transfection, media was changed to IGM and cells were infected with a low-defective stock of A/WSN/1933 at an MOI of 0.25. Infection was allowed to proceed for 72h, and the supernatant harvested, centrifuged at 300g for 4 minutes to remove cellular debris, and aliquoted into cryovials to be stored at -80^*°*^C. To sequence viral supernatant for Fig. 3, 100*µ*l of supernatant was mixed with 700 *µ*l of lysis buffer and purified using the Quick-RNA Miniprep Kit.

### Interferon selection experiments

For all experiments, we calculated the relative fraction of the library consisting of our indicated delVG versus full-length to identify appropriate multiplicities of infection. This was determined by using a delVG-specific qPCR and comparing this value to a qPCR that concurrently measures both full-length and delVG segments.^17^ Primers for these calculations provided in Supplementary Data 6. For PB1_177:385_ measurements, libraries were used to infect 30 million A549 *IFNL1* reporter cells seeded across three 15cm plates that were previously transduced to express PB1 at an MOI calculated on PB1_177:385_ of 0.2. We calculated our libraries comprised approximately 10% of the total library using qPCR, and so this was an infectious MOI of 2. For PB2_322:176_ and PA_328:137_ we did not use complementing cells, and as we calculated the majority of each population was our delVG as calculated by qPCR (*∼* 98% for PB2_322:176_, and *∼* 60% for PA_328:137_). To ensure sufficient complementation to allow delVG species to replicate we infected with our libraries at an MOI 1 as calculated for each delVG using qPCR, and added an additional wild-type helper virus at an MOI of 3, to infect a total of 30 million A549 *IFNL1* reporter cells across three T180 flasks. As performed in Mendes et al. (2021)^17^, we allowed infections to proceed for 14 hours prior to using magnetic enrichment to select for LNGFR-positive cells using MACSelect LNGFR microbeads (Miltenyi Biotec, # 130-091-330) using LS columns (Miltenyi Biotec, # 130-042-401). All LNGFR-enriched (*IFNL1* -enriched) cells were retained, and 5 million LNGFR-depleted (*IFNL1* -depleted) cells were retained for a matched analysis. RNA from both populations was purified using either the Monarch Total RNA Miniprep Kit (New England Biolabs, #T2010) for PB1_177:385_, or the Quick-RNA Miniprep Kit (Zymo Research Corp, Cat #R1055) for PB2_322:176_ and PA_328:137_.

### Preparation of sequencing libraries

Single-cell RNAseq datasets were generated using the 10x Genomics protocol using the Chromium Single Cell 3’ Library and Gel Bead kit v2, targeting 2000 cells for analysis with a 5% spike in of uninfected canine cells (MDCK-SIAT1) for inferences of contamination as previously-described, using a combined flu, canine, and human reference sequence.^26^

Sequencing of delVG variant libraries was largely performed as previously-described with minor modifications.^27^ Sequencing initial delVG plasmid libraries was performed using primers that sit outside of the mutated region to capture the entire delVG sequence, and overlapping amplicons such that each position was captured independently of primer binding. For sequencing viral RNA from delVG libraries we either used standard amplicon sequencing of vRNA, or, to resolve terminal nucleotides, 5’ RACE sequencing of vRNA and cRNA. For the former, cDNA was generated using Superscript III (Invitrogen, #18080400) with a with a universal vRNA primer with adapter from Hoffmann *et al*., according to the manufacturer’s protocol.^36^ For interferon selection PCR experiments, initial cDNA production used 100ng of RNA input, followed by qPCR to measure absolute delVG copies as compared between *IFNL1* -enriched and *IFNL1* -depleted samples. As *IFNL1* -depleted samples invariably had lower delVG content, initial RNA input was re-normalized to match *IFNL1* -enriched samples to ensure both were assessed at equivalent molecular depth. Supernatant samples simply used the maximum volume of RNA allowable in the reaction. cDNA was thereafter amplified with primers targeting the 5’ and 3’ termini, amplifying the entire molecule. Gel electrophoresis was used to separate out delVG amplicons from contaminating full-length PB2, PB1, or PA, followed by gel purification of the appropriately-sized band (New England Biolab’s Monarch DNA Gel Extraction Kit, #T1020). This material was then used for a set of targeted amplicons as in our plasmid sequencing, appending P5 and P7 partial adapters, gel-purified, and indexed for Illumina sequencing.

For 5’ RACE for delVG libraries, cDNA was generated using the Template Switching RT Enzyme Mix (New England Biolabs, #M0466) according to manufacturer’s protcol using either cRNA-specific or vRNA-specific primers with a custom template-switching oligonucleotide as previously described.^27,32,70^ For delVG libraries subsequent products were subject to a PCR spanning across the delVG junction and gel-purified to disambiguate delVG from full-length templates. This was followed by a PCR targeted the 5’ end of the vRNA, or cRNA, to add partial Illumina adapters for subsequent indexing PCR and sequening.

For 5’ RACE of high delVG populations as provided in Fig. 3a, a single product pertaining to both full-length and delVG variants of that segment was targeted for amplification, addition of partial Illumina adapters, and sequencing. For total RNAseq, RNA samples were provided to Azenta Genewiz for ribosomal RNA depletion and stranded RNAseq.

Primer sequences for all reactions provided and annotated in Supplemental Data 6.

All sequencing was performed on an Illumina NovaSeq platform using 2×150 paired-end reads.

### Computational analysis

To analyse single-cell RNAseq data from Fig. 1 we processed alignments from Cellranger 1.3.1 as described above. Reference for this and all other human alignments was GRCh38, reference for canine alignments was CanFam3.1, and a custom reference generated for A/WSN/1933 with a GTF corresponding to all major mRNAs observed in this virus was used for flu alignments. Due to relatively low contamination of transcripts of interest (flu and interferons), a strict cutoff was used of 1.5x the maximal number of UMIs observed in the canine cells, rather than the more nuanced assumption that this is a Poisson-mediated process. This was due to sparsity of sampling of extracellular reads, with insufficient data to infer an expected read distribution from contamination, and so we erred on the side of using a strict cutoff to exclude false positives.

Analysis of variant libraries was similar to that performed previously, but with several key modifications and improvements.^27^ First, as all amplicons were pooled for a given sample, we re-separated these pooled samples into individual, paired, FASTQ files. To do so, we enforced a perfect match of the first 15nt of each read (R1 and R2) to primers used to amplify a given amplicon. This excludes any off-target, aberrant, PCR products from consideration, allowing for better discrimination of per/site variant frequencies. For RACE samples, we next had to consider both caps as well as additional 5’ extensions such as a repeated template-switching oligonucleotide (TSO) or incorporation of cDNA primer. For this, we identified the spacing between the TSO and the putative end of the vRNA or cRNA. We permitted up to five mismatches throughout the entire TSO, and three mismatches in the 12 nucleotides used to identify the 5’ end of the vRNA or cRNA. If a sequence fails to meet those requirements it was excluded from further analysis. After identifying TSO and 5’ vRNA/cRNA locations, the space between the two was calculated, if this space was three or less, consistent with the average number of untemplated nucleotides added by reverse transcriptase during template-switching reactions, we retained this sequence as uncapped.^32^ If greater than 10, this sequence was retained as capped in our code, however was excluded from analyses here which focus on vRNA or cRNA and not mRNA. If neither, these sequences were excluded. Untemplated nucleotides were then trimmed to provide better alignment metrics and resolve nucleotide variance at the 5’ end of the cRNA/vRNA. The same matching schema were used to process reads from our plasmid library such that the enforced matching does not produce inappropriately reduced measurements of variance in RNA samples versus our original library, as described previously.^27^ After processing all FASTQ files, we next mapped each to the specific amplicon from which they were derived using STAR.^71^ We enforced end-to-end mapping, excluded splicing using alignIntronMax 1 and a custom GTF consisting of just the amplicon, and enforced an average mismatch rate of <2% to exclude aberrant amplicons from further consideration. Per/site nucleotide identity was then assessed using a custom script, accounting for paired-ends and taking the better quality score call when in conflict, using a q-value cutoff of 30. For each individual amplicon, sites on which our primers sat were excluded from per/site nucleotide analyses. Indels are provided in supplement but were not considered in our work here.

For 5’ RACE of previously-isolated high delVG populations sorted by interferon as presented in Fig. 3a, we used our custom script described above to identify capped species, and trim untemplated nucleotides, and then aligned only our read proximal to the 5’ end (R2) to a STAR reference generated from the complete A/WSN/1933 sequence. We then used a custom script to infer per/site nucleotide identity from this single-end mapped data, with a quality score cutoff of 30 as above. Indels are provided in supplement but were not considered in our work here.

To analyse stranded RNAseq from Fig. 4a,b, we first used Trimmomatic with the following parameters: 2 seed mismatches, a palindrome clip threshold of 30, a simple clip threshold of 10, a minimum adapter length of 2, keep both reads, a lead of 20, a sliding window from 4 to 15, and a minimum retained length of 36.^72^ Reads were then mapped against a combined human flu reference using STAR, using PB1_445:350_ instead of full-length PB1 and using strand-specific GTF for influenza. STAR parameters used default parameters, unlike above. HT-seq count was then used to produce gene counts, using the reverse flag to account for R2 mapping in the (+)-sense rather than R1; this is the standard orientation for Illumina stranded library preparation.^38^ Spliced transcripts were collapsed due to large degree of ambiguity in assignment, however assessment of bamfiles through IGV did reveal the vast majority of M and NS-derived mRNAs were unspliced, as expected. Counts were then analysed by DEseq2 using a Python implementation.^37,73^ As (+)-sense reads were nearly completely suppressed by baloxavir, and previous literature established little to no cRNA is produced in the absence of *de novo* polymerase production, all (+)-sense reads were assigned to mRNA. For Fig. 4b, counts were corrected for length of each individual segment with an assumption that fragment frequency is directly proportional to length before being summed and used to calculate frequency of mRNA or vRNA derived from each segment. For assessment of polyA tails, we used the raw fastQ files rather than alignment, due to ambiguity and missing alignments of polyadenylated species. For both R1 and R2, separately, we used the sequences 5’-CATGAAGGACAAGCTAAATT’-3’ and 5’-AATTTAGCTTGTCCTTCATG-3’ for R1 and R2, respectively, to pull trimmomatic-trimmed reads containing the 3’ sequence adjacent to the polyuracil tract in (+)-sense RNA only. Concurrently, using grep with regular expression, we pulled 9nt including and proceeding the polyuracil tract in the (+)-sense orientation. For the 3nt after the polyuracil tract, we manually assessed whether they were consistent with polyadenylation (all A’s as read in the (+)-sense orientation) or ccRNA/cRNA (standard 3nt derived from flu cRNA). Each read was assessed independently given sparsity of coverage. For assessment of mutations at the U16/U20 positions, in Supplementary Fig. 7, we pulled only (-) sense, PB1_445:350_ vRNA, reads from an aligned bamfile using samtools and assessed the per/site nucleotide identity using the same framework and script as for our delVG libararies.^74^

The following software and package versions were used: cellranger 1.3.1, Python 3.12.2, Trimmomatic 0.39, STAR 2.7.11b, Samtools 1.19, FastQC 0.12.1, numpy 1.26.4, matplotlib 3.8.3, seaborn 0.13.2, pandas 2.2.1, scipy 1.12.0, and statsmodels 0.14.1.^71,72,75–81^

All code was run on the San Diego supercomputer center, Triton shared computing cluster.^82^

## Data Availability

The processed files and raw sequencing data generated in this study have been deposited in the NCBI gene expression omnibus under accessions GSE276697, GSE350335 and GSE350336.

## Code Availability

The analysis code used in this study is provided at https://github.com/a5russell/IAV_delVG_polyuracil/.

## Acknowledgments

All work described in this manuscript except for partial salary of SNZH was funded by the National Institute of General Medicine (NIGMS) of the NIH under grant R35GM147031 awarded to ABR. Work performed by SNZJ was in part supported by the NIGMS of the NIH under Grant T32GM133351. The content is solely the responsibility of the authors and does not necessarily represent the official views of the National Institutes of Health. We thank Andrew Mehle for the kind gift of MDCK cells expressing PB2. We thank the Russell lab for insightful discussion in the preparation of this manuscript. The writing of this manuscript, literature analysis, and computational analysis were performed without the aid of LLMs.

## Author Contributions Statement

ABR, MM, SNZJ, designed the study, performed experiments, and analysed the data. ABR and SNZJ prepared the manuscript.

## Supplementary Figures, Tables, and Files

**Supplementary Figure 1.**
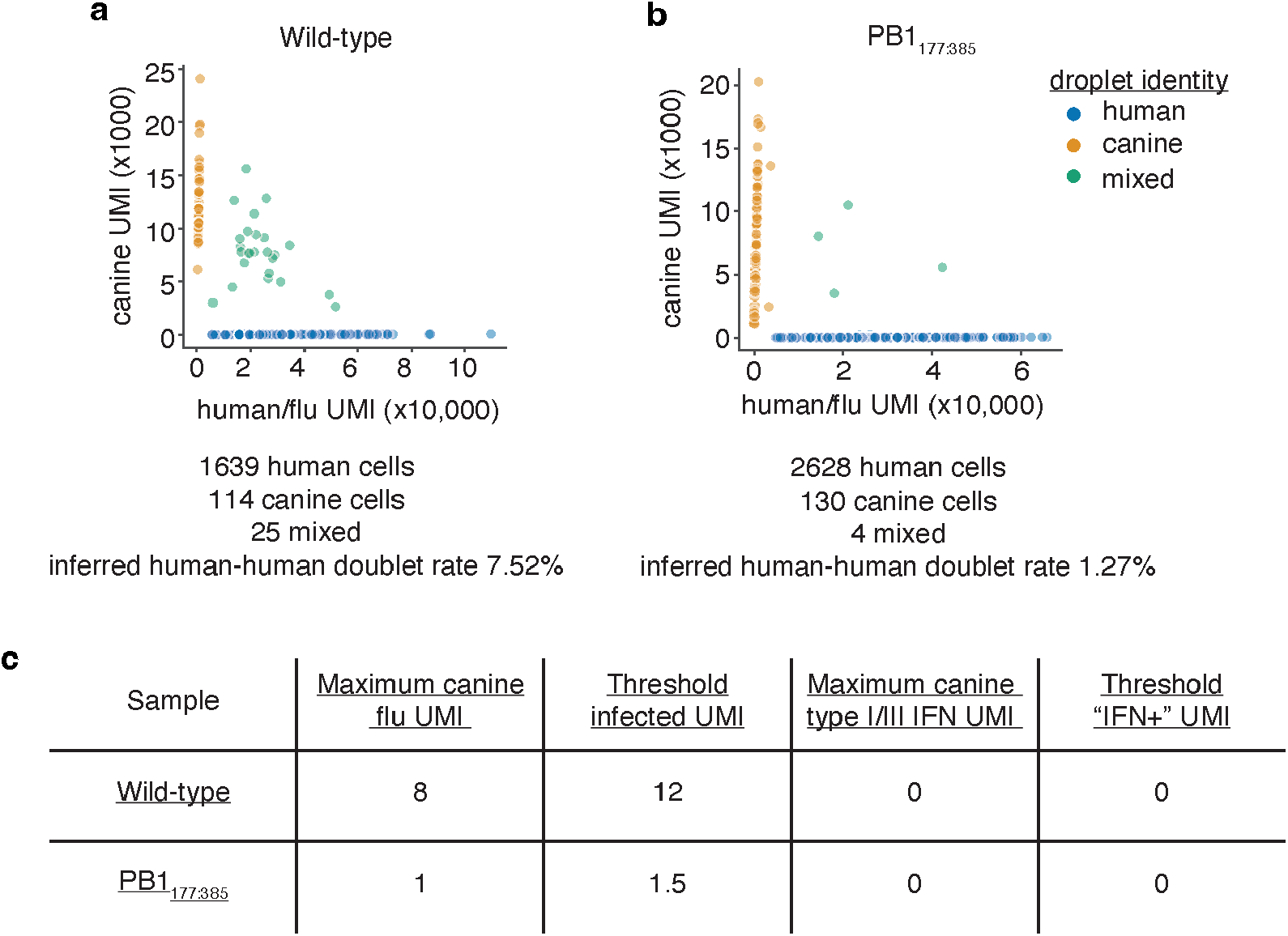
Single-cell RNAseq metrics from data presented in Fig. 1. **a**,**b** Annotation of A549 (human) cells infected with indicated strains at an MOI of 0.1 mixed with an uninfected MDCK-SIAT1 (canine) cell line for multiplet and extracellular RNA measurements. **c** UMI thresholds set on the maximal UMI observed in canine cells. While conservative, this allows a strict, empiric, threshold to be set. Multiplet and extracellular contamination analysis was performed as previous.^16^

**Supplementary Figure 2.**
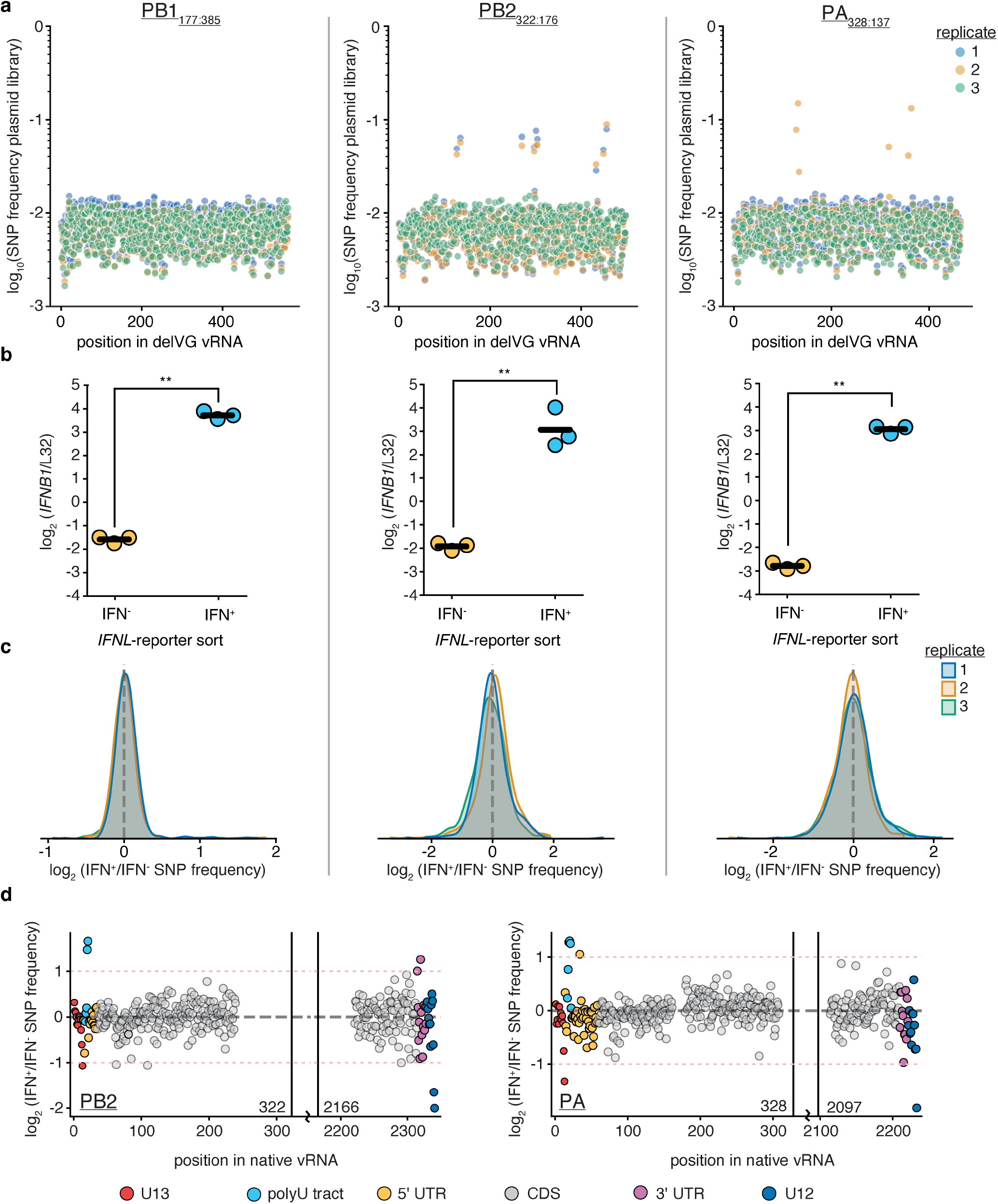
Library metrics for Fig. 2. **a** Per site SNP frequences per replicate per library generated by mutagenic PCR. **b** A549 *IFNL1* reporter cells were infected at an MOI calculated on delVG libraries alone of 0.2 (PB1) or 1 (PB2, PA). For PB1 cells were transduced to express PB1, for PB2 and PA complementing virus was added to a final MOI of 3. At 14hpi cells were sorted using a magnetically-enrichable *IFNL1* reporter. Enrichment assessed here by qPCR. n=3, asterisks indicate significant difference, two-tailed t-test p<0.05. **c** Distribution of SNP frequency ratios as calcuated in Fig. 2a. To identify significant sites we assessed each site against the population background, which we demonstrate here follows a predictable distribution centered at no-effect (0, dottted line). **d** All sequenced sites for PB2_322:176_ and PA_328:137_ presented as in Fig. 2a.

**Supplementary Figure 3.**
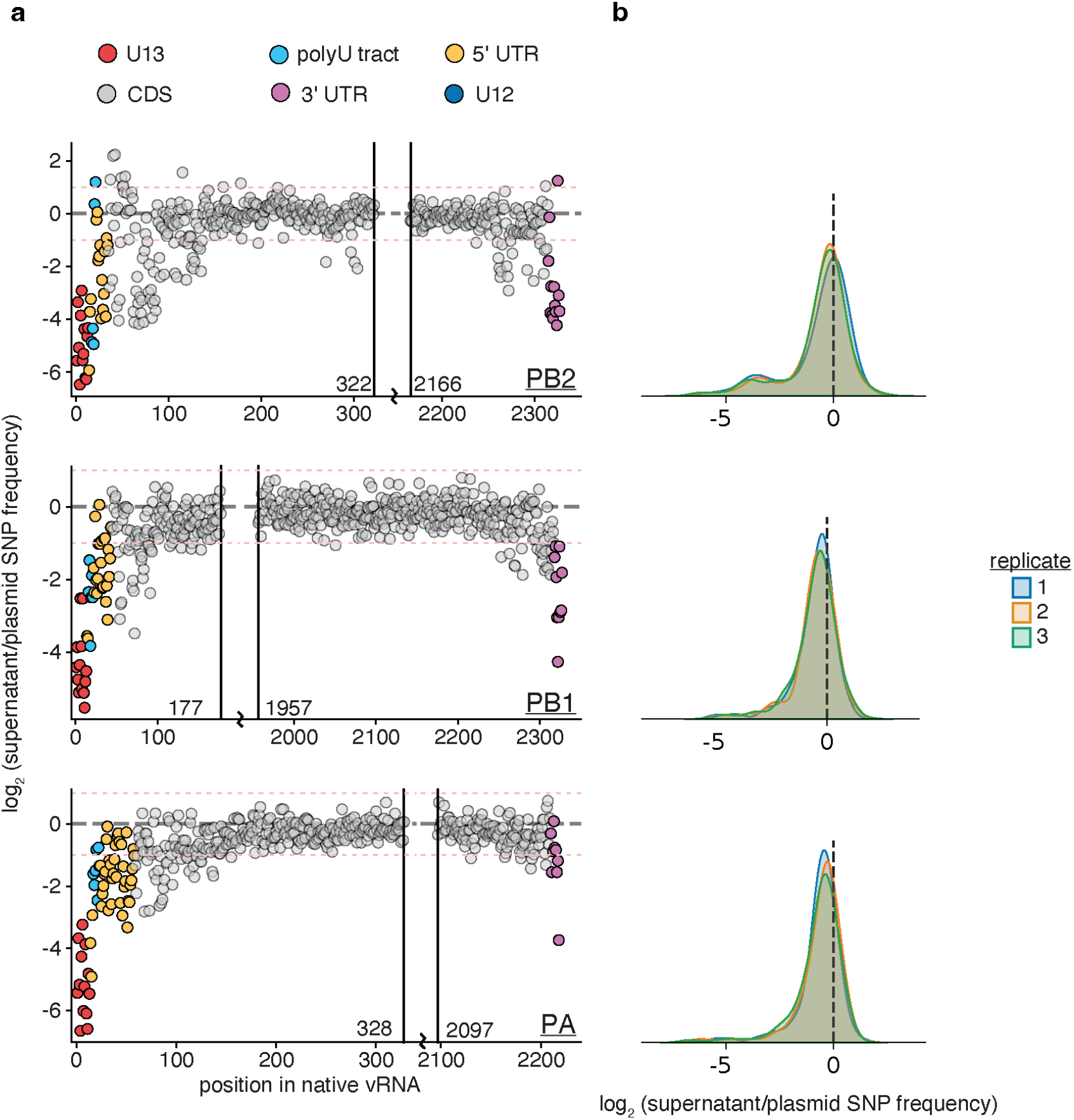
Library metrics describing selection measured by progression of libraries to viral supernatant. **a** Mean ratio of non-wild-type nucleotide frequency at each position in each library comparing supernatant virion RNA after reverse genetics rescue with complemetning virus versus the original plasmid library. Red dotted lines indicate two-fold differences in enrichment or depletion. Coordinates provided respective to full-length vRNA, regions of known functional significance noted. U12 and adjacent nucleotides excluded as there is no reliable cRNA measurement from supernatant virions, and we inferred ends using 5’ RACE. **b** Distribution of effects from (a), revealing a leftward skew consistent with a large number of negative-effect mutations. As such analysis in Fig. 3b was performed relative to no effect, rather than against the population average.

**Supplementary Figure 4.**
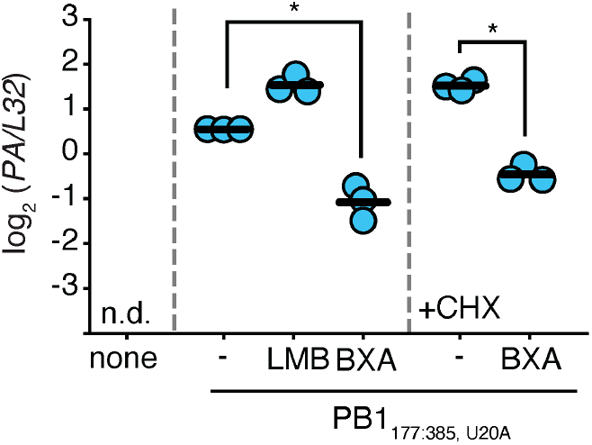
Influence of inhibitors on viral transcription. Experiment from Fig. 4c measuring levels of the flu RNA *PA* versus the housekeeping control *L32*. n =3. Asterisks indicate significant reduction of *PA* within panels against no leptomycin (LMB) or baloxavir acid (BXA), one-tailed t-test with Benjamini-Hochberg correction, FDR<0.05.

**Supplementary Figure 5.**
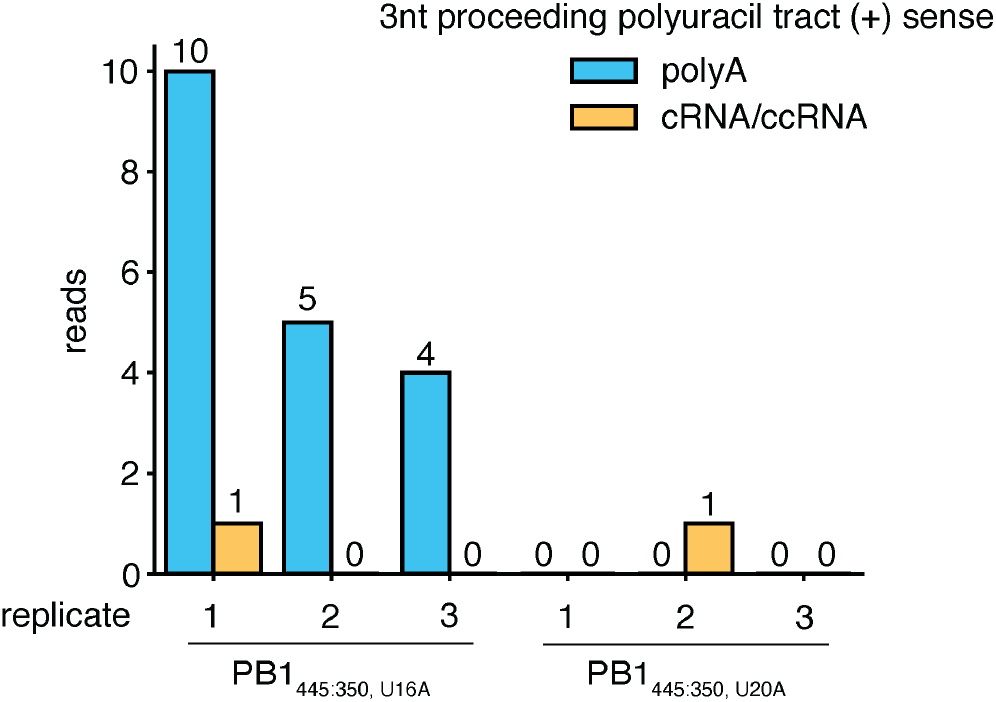
Rare detection of polyadenylation state from Fig. 4. Experiment from Fig. 4a. As mapping is unreliable around the polyadenylation sequence, reads (forward or reverse) matching 20nt 5’ of the PB1_445:350_ mRNA (+)-sense polyadenylation site were recovered and the adjacent 9nt 3’ of this sequence were analysed as either supporting polyadenylation (all A’s), supporting cRNA, or ccRNA (3nt that proceed the standard polyuracil tract, including introduced U16A or U20A mutations).

**Supplementary Figure 6.**
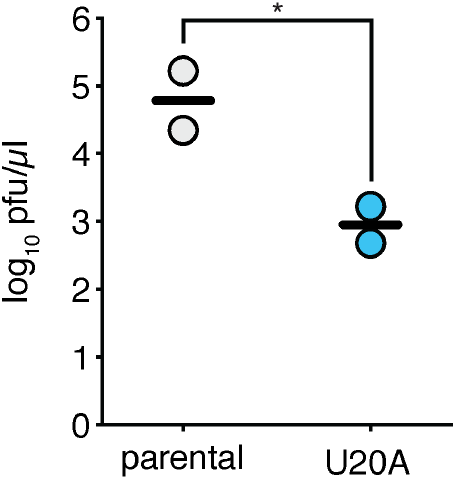
U20A mutation decreases viral titers for a full-length PB1 template. Titers of full-length PB1 with a start codon mutation and the indicated variants after reverse genetics as described in Methods. Asterisk indicates significant difference, two-tailed t-test, p<0.05, n=2.

**Supplementary Figure 7.**
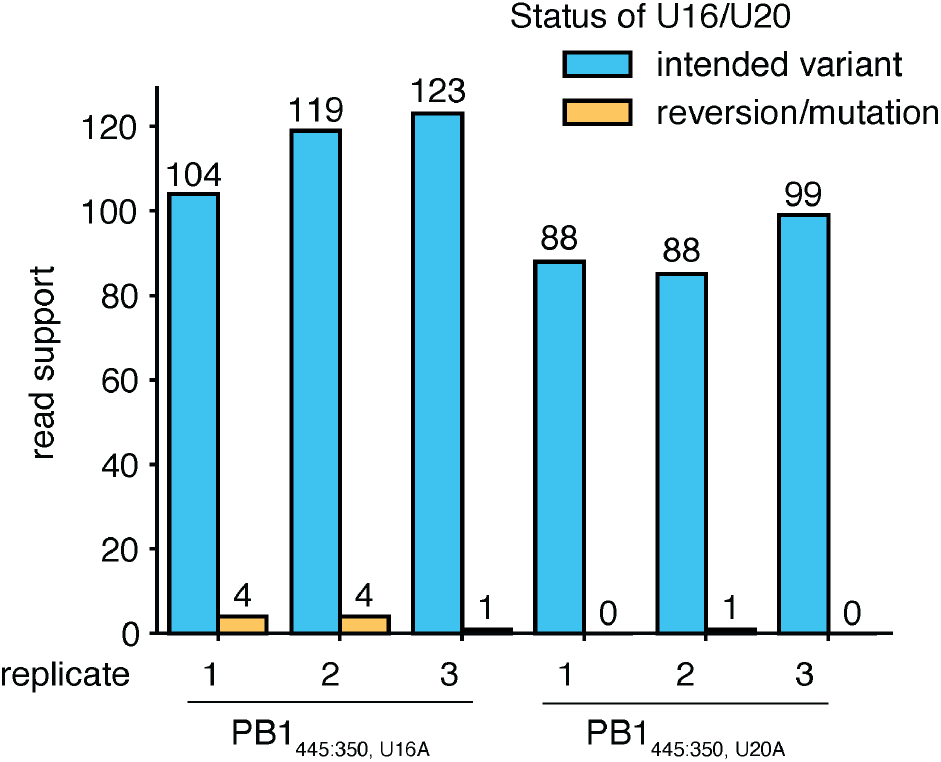
U20A and U16A mutations are retained. Read support for the intended U16A or U20A mutations from (-) sense RNA reads only mapped to PB1_445:350_ from Fig. 4a. Positions inferred from paired-end data using standard quality score cutoff of 30. Bases called at all positions provided in Supplementary Data 5.

**Supplementary Data 1. Genome sequence of A/WSN/1933 and variants**.

Genome sequences of A/WSN/1933 and variants used in this study.

**Supplementary Data 2. delVG junctions**.

delVG junctions identified in scRNAseq data using a UMI cutoff of two and only considering cells positively called as infected. delVG meeting this cutoff were only identified in wild-type infection. Coordinates reflect (+) sense boundaries of junctions with last base retained at 5’ and 3’ ends.

**Supplementary Data 3. ISGs**.

List of ISGs derived from Stevens *et al*.(2022)^23^ using cutoffs of greater than a 2-fold increase at a FDR of 0.1 upon interferon treatment.

**Supplementary Data 4. Natural delVG variation**.

Per/site nucleotide identity from interferon-enriched and -depleted populations with targeted sequencing of the 5’ vRNA of PB2, PB1, and PA from Mendes and Russell 2021.^17^

**Supplementary Data 5. Lack of variation in U16A and U20A positions**.

Per/site nucleotide identity in (-) sense RNA from PB1_177:385_ from total RNAseq presented Supplementary Fig. 7.

**Supplementary Data 6. Primer list**.

Sequence of all primers used in this study.

